# CRISPR/Cas9 knock-in strategy to evaluate phospho-regulation of SAMHD1

**DOI:** 10.1101/2022.01.04.475018

**Authors:** Moritz Schüssler, Paula Rauch, Kerstin Schott, Adrian Oo, Nina Verena Fuchs, Baek Kim, Renate König

**Author notes:** Address correspondence to Renate König.

## Abstract

Sterile α motif (SAM) and HD domain-containing protein 1 (SAMHD1) is a potent restriction factor for immunodeficiency virus 1 (HIV-1), active in myeloid and resting CD4^+^ T cells. As a dNTP triphosphate triphosphohydrolase (dNTPase), SAMHD1 is proposed to limit cellular dNTP levels correlating with inhibition of HIV-1 reverse transcription. The anti-viral activity of SAMHD1 is regulated by dephosphorylation of the residue T592. However, the impact of T592 phosphorylation on dNTPase activity is still under debate. Whether additional cellular functions of SAMHD1 impact anti-viral restriction is also not completely understood.

We use BlaER1 cells as a novel human macrophage transdifferentiation model combined with CRISPR/Cas9 knock-in (KI) to study SAMHD1 mutations in a physiological context. Transdifferentiated BlaER1 cells, resembling primary human macrophages, harbor active dephosphorylated SAMHD1 that blocks HIV-1 reporter virus infection. Co-delivery of Vpx or CRISPR/Cas9-mediated SAMHD1 knock-out relieves the block to HIV-1. Using CRISPR/Cas9-mediated homologous recombination, we introduced specific mutations into the genomic *SAMHD1* locus. Homozygous T592E mutation, but not T592A, leads to loss of HIV-1 restriction, confirming the role of T592 dephosphorylation in the regulation of anti-viral activity. However, T592E KI cells retain wild type dNTP levels, suggesting the antiviral state might not only rely on dNTP depletion.

In conclusion, the role of the T592 phospho-site for anti-viral restriction was confirmed in an endogenous physiological context. Importantly, loss of restriction in T592E mutant cells does not correlate with increased dNTP levels, indicating that the regulation of anti-viral and dNTPase activity of SAMHD1 might be uncoupled.

**Importance:** Sterile α motif (SAM) and HD domain-containing protein 1 (SAMHD1) is a potent anti-viral restriction factor, active against a broad range of DNA viruses and retroviruses. In myeloid and resting CD4^+^ T cells, SAMHD1 blocks reverse transcription of immunodeficiency virus 1 (HIV-1), not only inhibiting viral replication in these cell types, but also limiting the availability of reverse transcription products for innate sensing of HIV-1. Manipulating SAMHD1 activity could be an attractive approach to improve HIV-1 therapy or vaccination strategies. Anti-viral activity is strictly dependent on dephosphorylation of SAMHD1 residue T592, however the mechanistic consequence of T592 phosphorylation is still unclear. Here, we use BlaER1 cells as an alternative myeloid cell model in combination with CRISPR/Cas9-mediated KI to study the influence of SAMHD1 T592 phosphorylation on anti-viral restriction and control of cellular dNTP levels in an endogenous context. By using this novel approach, we were able to genetically uncouple SAMHD1’s anti-viral and dNTPase activity with regard to regulation by T592 phosphorylation. This suggests that SAMHD1 dNTPase activity may not exclusively be responsible for the anti-lentiviral activity of SAMHD1 in myeloid cells. In addition, our toolkit may inspire further genetic analysis and investigation of SAMHD1-mediated restriction, as wells as its cellular function and regulation, leading to a deeper understanding of SAMHD1 and HIV-1 biology.

## Introduction

Sterile α motif (SAM) and HD domain-containing protein 1 (SAMHD1) is a potent anti-viral restriction factor with broad anti-viral activity against a number of viruses, among them lenti- and non-lenti retroviruses (for review see (1)). In particular, HIV-1 replication is restricted in myeloid cells and resting CD4^+^ T cells (2–6). SAMHD1 depletion leads to an increase in intermediates of reverse transcription (RT), especially late cDNA products, indicating that SAMHD1 inhibits the RT process (3, 7, 8).

SAMHD1 is a cellular dNTP triphosphate triphosphohydrolase (dNTPase). It is active as a tetramer, regulated by binding of GTP/dGTP and dNTPs to primary and secondary allosteric sites, respectively (9). Therefore, the obvious assumption might be that SAMHD1 inhibits HIV-1 replication through depletion of dNTPs, the substrate for HIV-1 RT (10, 11). Providing exogenous desoxyribonucleotides (dNs) rescues HIV-1 replication in cells expressing SAMHD1 (5, 11). In addition, SAMHD1 mutants shown to lack dNTPase activity, both *in vitro* or in cells, lose their restrictive potential, when overexpressed in phorbol 12-myristate-13-acetate (PMA)-activated macrophage-like U937 cells (2, 11–13). However, SAMHD1 dNTPase activity might not be sufficient for HIV-1 restriction (1, 14). It is hypothesized that additional SAMHD1 mediated functions like modulation of immune signaling, resolution of stalled replication forks and R-loops, RNA binding, or its role in DNA damage response, might contribute to the restrictive phenotype (1, 15–18).

Only SAMHD1 dephosphorylated at residue T592 is active against HIV-1 (19–21). SAMHD1 is phosphorylated in cycling cells by cyclin dependent kinases CDK1 and CDK2 in complex with Cyclin A2 in S and G_2_/M phase (20, 19). At mitotic exit, SAMHD1 is rapidly dephosphorylated at residue T592 due to the action of the PP2A-B55α phosphatase complex (22). While the effect of SAMHD1 T592 phosphorylation on HIV-1 restriction is consistently demonstrated, the consequence for its dNTPase activity is still under debate. Biochemical approaches to measure the effect of SAMHD1 T592 phosphorylation and phosphomimetic mutants on SAMHD1 tetramer formation and dNTPase activity have not been able to reveal a functional relationship (12, 20, 21, 23, 24). Still, cell cycle dependent SAMHD1 phosphorylation, loss of HIV-1 restriction and increased dNTP levels in S- and G_2_/M phase in synchronized HeLa cells show a clear timely correlation (22). In contrast, mutagenic analysis of T592 site in myeloid cells challenges a causative link. Phosphoablative T592A or T592V, but not phosphomimetic T592E or T592D mutants, were able to inhibit HIV-1 replication, when overexpressed in PMA activated U937 cells (12, 20, 21, 25). Conversely, not only phosphoablative but also phosphomimetic SAMHD1 T592 mutants efficiently limited the cellular dNTP pool (14, 20, 21). This obvious discrepancy might be due to biological reasons (for details refer to discussion and review (1)). However, also technical limitations might be the cause for this problem.

Genetic studies of SAMHD1 phospho-mutants in myeloid cells are currently limited to PMA-activated macrophage-like THP-1 or U937 cells. As treatment with PMA can activate non-physiological intracellular pathways (26, 27), alternative myeloid models are needed, which ideally be both genetically amendable and based on physiological myeloid differentiation pathways.

So far, anti-viral restriction has been tested with mutant constructs of SAMHD1 using lenti- or retroviral transduction. In this case, an exogenous promotor mediates overexpression of SAMHD1. The use of CRISPR/Cas9 allowed us to modify *SAMHD1* within the native genetic environment and to analyze the impact of selected mutations on anti-viral restriction in a physiological context, avoiding potential unwanted effects of mutant protein overexpression.

Here, we use CRISPR/Cas9-mediated knock-in (KI) in combination with transdifferentiated macrophage-like BlaER1 cells as a tool to study the impact of SAMHD1 T592 phosphorylation on HIV-1 restriction and dNTP pools in myeloid cells. Transdifferentiated macrophage-like BlaER1 cells expressed SAMHD1, which was dephosphorylated at residue T592. Concomitantly, transdifferentiated BlaER1 cells restricted HIV-1 replication in a SAMHD1 dependent manner. Introduction of SAMHD1 homozygous T592E mutations via CRISPR/Cas9 KI led to loss of HIV-1 restriction, while SAMHD1 T592A mutants maintained their anti-viral activity. Remarkably, neither endogenous SAMHD1 T592E, nor T592A mutants, had an impact on cellular dNTP levels in transdifferentiated BlaER1 cells, indicating that the regulation of anti-viral and dNTPase activity of SAMHD1 might be uncoupled.

## Results

### Transdifferentiated BlaER1 cells express SAMHD1 dephosphorylated at residue T592

Myeloid models to study HIV-1 restriction by mutagenesis are very limited. Transdifferentiated BlaER1 cells are a novel myeloid cell model, which has successfully been used to study innate immune signaling in macrophage-like cells (28, 29). Here, the native, B-lineage derived BlaER1 cells undergo macrophage transdifferentiation by induction of the myeloid transcription factor C/EBPα. In order to test whether these cells can serve as a model to study SAMHD1 mediated anti-viral restriction, we analyzed SAMHD1 expression in transdifferentiated BlaER1 cells. Flow cytometry analysis of transdifferentiated BlaER1 cells showed loss of B cell marker CD19 and acquisition of surface expression of the macrophage marker CD11b (Fig. 1A), as demonstrated earlier (29). Transdifferentiation of BlaER1 cells using an adopted protocol, was highly reproducible and yielded 89.3 ± 8.8% (n = 33) of CD19^−^ CD11b^+^ cells in living BlaER1 cells expressing GFP (Fig. 1B). Transdifferentiated BlaER1 cells expressed levels of SAMHD1 comparable to cycling THP-1 cells (Fig. 1C), whereas native BlaER1 cells show no SAMHD1 expression. As T592 phosphorylation in SAMHD1 is the major regulator of antiviral restriction (19, 20, 22), we analyzed the phosphorylation status in transdifferentiated BlaER1 cells. Relative SAMHD1 T592 phosphorylation was 31-fold lower in transdifferentiated BlaER1 compared to cycling THP-1 cells (0.032 ± 0.013 relative SAMHD1 pT592 normalized to cycling THP-1, n = 6) and in fact was barely detectable by immunoblotting even after long exposure times (Fig. 1C). Absence of SAMHD1 pT592 correlated well with the reported G1/G0 cell cycle arrest in transdifferentiated BlaER1 cells (22, 29), as well as with low cyclin A2 expression (Fig. 1C), which in complex with CDK1 and CDK2 is known to mediate T592 phosphorylation (19). Thus, macrophage-like transdifferentiated BlaER1 cells expressed dephosphorylated SAMHD1 at residue T592, suggesting it to be anti-virally active.

**Figure 1:**
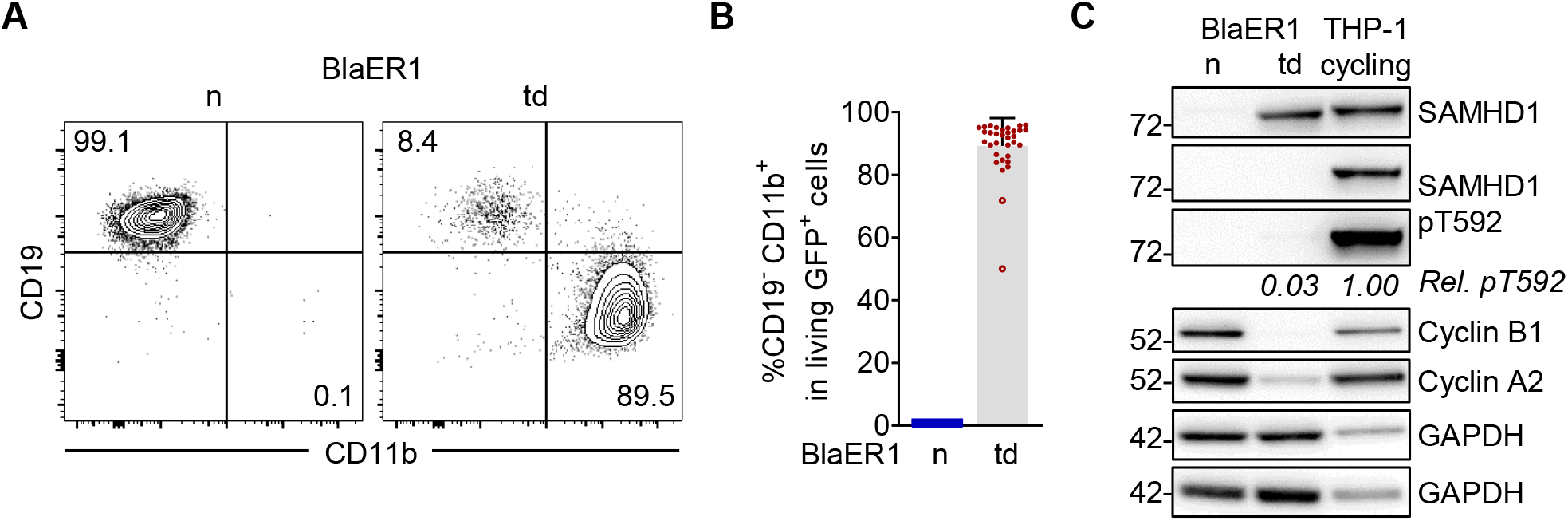
Transdifferentiated BlaER1 cells express SAMHD1 which is dephosphorylated at residue T592. **(A)** Representative flow cytometry analysis of CD19 and CD11b surface expression in native (n) and transdifferentiated (td) BlaER1 cells. Relative frequencies of CD19^+^ CD11b and CD19 CD11b^+^ cell populations are indicated as % of living GFP^+^ cells (n = 33). **(B)** Relative quantification of macrophage-like CD19 CD11b^+^ cells in living GFP^+^ native (n) or transdifferentiated (td) BlaER1 cells. Every dot represents an individual transdifferentiation approach. Experiments in which transdifferentiated BlaER1 cells show < 75% CD19 CD11b^+^ cells in living GFP^+^ cells were excluded from downstream analysis (open circles). Error bars represent standard deviation (n_n_ = 30, n_td_ = 33). **(C)** Representative immunoblot analysis of SAMHD1, Cyclin B1 and Cyclin A2 expression in native (n) and transdifferentiated (td) BlaER1 cells, as well as cycling THP-1 cells. GAPDH serves as a loading control. Mean signal of SAMHD1 T592 phosphorylation (pT592) relative to total SAMHD1 expression in transdifferentiated BlaER1 cells was normalized to cycling THP-1 (n = 6).

### SAMHD1 is a major restriction factor against HIV-1 in transdifferentiated BlaER1 cells

To define the restrictive capacity of SAMHD1 in the context of transdifferentiated BlaER1 cells, we infected the cells with a single-cycle HIV-1 luciferase reporter virus (HIV-1-luc), in presence or absence of virus like particles containing Vpx (VLP-Vpx). VLP-Vpx treatment led to efficient degradation of SAMHD1 (0.013 ± 0.007 relative SAMHD1 expression normalized to no VLP-Vpx, n = 3) (Fig. 2A) and increased HIV-1-luc infection. Linear regression revealed a significant (*p = 0.0125*, n = 3, unpaired t-test) increase over a wide range of MOIs (Fig. 2B). To validate this further, we generated SAMHD1 knock-out (KO) BlaER1 cells using CRISPR/Cas9 ribonucleoprotein (RNP). Three independent SAMHD1 KO BlaER1 single cell clones were analyzed in detail and showed bi-allelic InDels at the intended target site (Fig. 2C), leading to a frameshift, the introduction of premature stop codons and therefore, absence of SAMHD1 expression in transdifferentiated BlaER1 cells (Fig. 2D). While SAMHD1 KO did not affect BlaER1 transdifferentiation (Fig. 2E), it strongly increased HIV-1-luc infection at 24 hpi, as compared to wild type (WT) cells. Significance of differences in the slopes of linear regressions are suggesting SAMHD1 to be a major restriction factor in these cells over a wide range of MOIs (*p < 0.0001* for Clone #1, 2 and 3, n = 7, One-way ANOWA) (Fig. 2F). In order to rule out a potential confounding effect of a minor CD11b^−^ native-like population, we developed a flow cytometry workflow combining the use of a single-cycle HIV-1 mCherry (HIV-1-mCherry) reporter virus together with staining for living CD11b^+^ cells. Thereby, we could specifically analyze infection in transdifferentiated CD11b^+^ macrophage-like BlaER1 cells. HIV-1-mCherry infection, as measured by %mCherry^+^ cells in CD11b^+^ living GFP^+^ transdifferentiated BlaER1 cells, was strongly increased upon SAMHD1 KO at 24 hpi (Clone #1 *p = 0.3633*, #2 *p = 0.0360*, #3 *p = 0.0013*, n = 5, Kruskal–Wallis test) (Fig. 2G and H). This indicates that SAMHD1 is a major anti-lentiviral restriction factor in macrophage-like transdifferentiated BlaER1 cells. We therefore conclude that transdifferentiated BlaER1 cells are an ideal model to study SAMHD1 mediated HIV-1 restriction.

**Figure 2:**
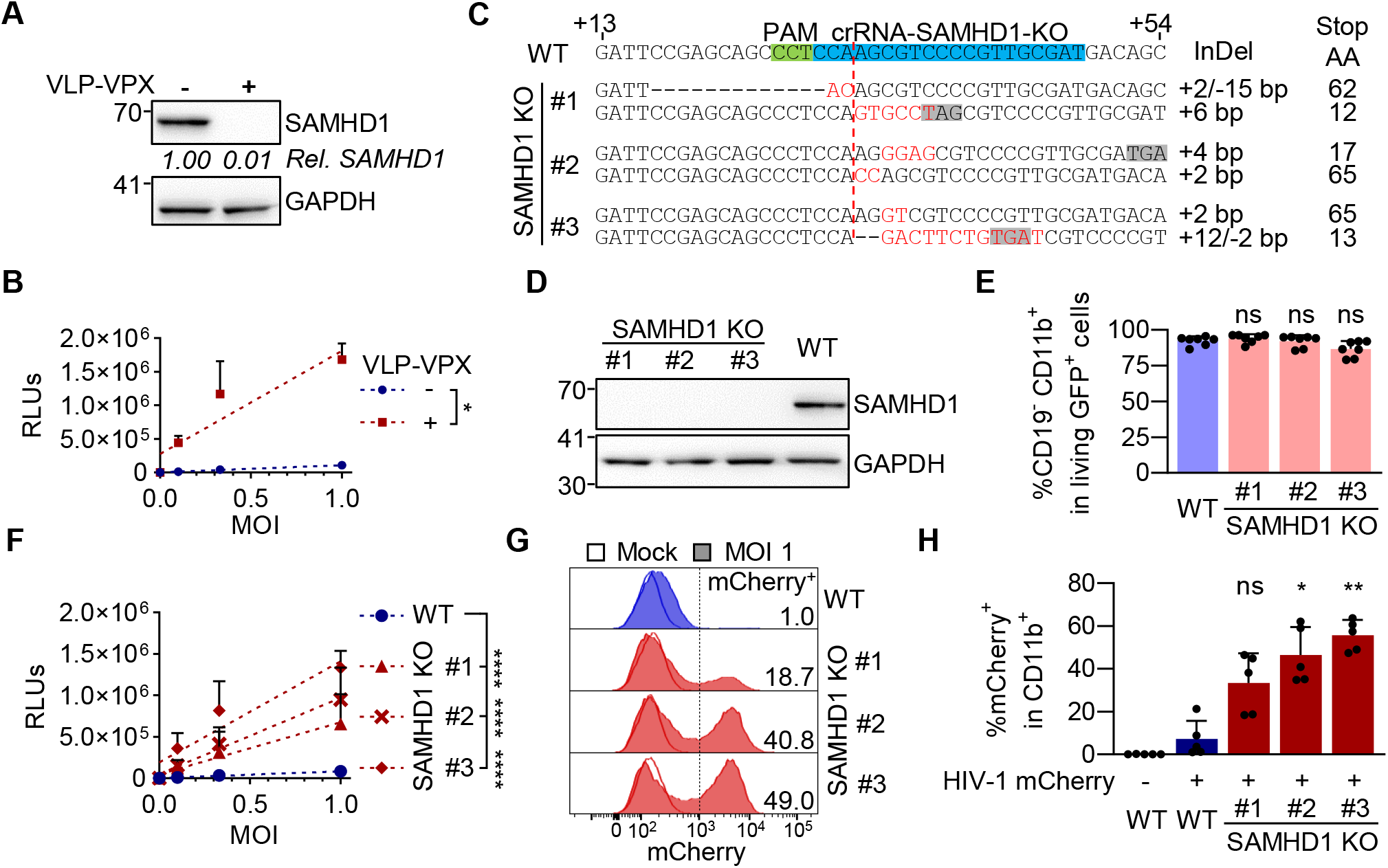
SAMHD1 restricts HIV-1 replication in transdifferentiated BlaER1 cells. **(A)** Transdifferentiated BlaER1 cells were treated with VLP-Vpx or medium for 24 h. SAMHD1 degradation was measured by immunoblot and quantified relative to GAPDH expression, followed by normalization to medium treated control (mean of n = 3). **(B)** VLP-Vpx or medium treated transdifferentiated BlaER1 cells were infected with VSV-G pseudotyped HIV-1 single-cycle luciferase reporter virus pNL4.3 E R luc at an MOI of 0.1, 0.33 and 1. Relative light units (RLUs) were quantified by luciferase measurement at 24 hpi. Linear regressions (dashed lines) were calculated and differences of slopes were tested for significance (n = 3, t-test). **(C)** BlaER1 cells were treated with CRISPR/Cas9 protein complexed with crRNA-SAMHD1-KO. Single cell clones were Sanger sequenced after TA-cloning to separate alleles and aligned to WT sequence. Insertions (red) and/or deletions (InDel) are indicated, as well as the position of the premature stop codon (gray), introduced by the respective genetic modification. **(D)** Genetically confirmed SAMHD1 knock-out (KO) clones were analyzed via immunoblot for SAMHD1 expression in transdifferentiated BlaER1 cells. GAPDH was used as loading control (n = 7). **(E)** Percentages of CD19 CD11b^+^ cells in living GFP_+_ transdifferentiated WT and SAMHD1 KO cells were quantified by flow cytometry (n = 7, One-way ANOWA). **(F)** RLUs in transdifferentiated BlaER1 WT and KO cell clones were quantified 24 hpi with pNL4.3 E R luc (VSV-G). Statistical significance of differences between linear regressions (dashed lines) in SAMHD1 KO clones compared to WT are indicated (n = 7, One-way ANOWA). **(G, H)** Transdifferentiated WT and SAMHD1 KO cell clones were infected with VSV-G pseudotyped HIV-1 single cycle mCherry reporter virus pNL4.3 IRES mcherry E^−^ R^+^ at MOI 1. Percentage of mCherry^+^ cells was quantified by flow cytometry in living GFP^+^ CD11b^+^ BlaER1 cells 24 hpi. **(G)** Representative histograms are shown for mock and HIV-1 mCherry_+_ reporter virus infected cells. Percentage of mCherry in living GFP^+^ CD11b^+^ BlaER1 cells is indicated. **(H)** Bar graphs indicate mean of experiments, dots individual biological replicates (n = 5, Kruskal–Wallis test). **(B, E, F, H)** Error bars correspond to standard deviation (* p < 0.05; ** p < 0.01; *** p < 0.001; **** p < 0.0001; ns, not significant).

### A pipeline to generate mutants of SAMHD1 by CRISPR/Cas9 mediated knock-in

So far, mutagenic analysis of SAMHD1 has been limited to model systems in which SAMHD1 is overexpressed by transient transfection or retroviral transduction. Overexpression of SAMHD1, especially in the context of phosphomimetic T592E or phosphoablative T592A mutation and their effect on viral restriction and intracellular dNTP levels, might affect functional readouts due to non-physiological expression levels, abnormal genomic context and altered post-translational regulation (1). To overcome this challenge, we decided to introduce SAMHD1 point mutations directly into the *SAMHD1* gene locus by CRISPR/Cas9 KI. Therefore, we developed a pipeline based on the introduction of RNPs and single-stranded DNA correction templates by electroporation, followed by an allele-specific PCR (KASP-genotyping assay screening) and rigorous validation by Sanger sequencing and quantitative genomic PCR to exclude large genomic deletions (qgPCR) (30) (Fig. 3A). We could identify single cell clones, displaying homozygous introduction of T592A and T592E mutations into the *SAMHD1* locus of BlaER1 cells (Fig. 3B). Quantification of allele numbers of SAMHD1 exon 16 revealed that the majority of homozygous single cell T592A and T592E KI clones still contained two alleles of SAMHD1 exon16 (Fig. 3C). However, we could identify one out of 8 clones analyzed (Clone X), which showed loss of one allele in qgPCR, indicative of pseudo-homozygosity (30). In total, we were able to generate and validate two homozygous T592A, as well as three homozygous T592E BlaER1 KI mutants, corresponding to a homozygous KI frequency of ~1% and highlighting the necessity of KASP-screening to reduce the number of KI candidates (Fig. 3D). Expression of SAMHD1 mutants in transdifferentiated T592A or T592E KI BlaER1 single cell clones is at similar level compared to WT protein in the respective parental cell line (Fig. 3E). SAMHD1 KI had no negative impact on BlaER1 transdifferentiation (Fig. 3F). In summary, using our pipeline we were able to introduce homozygous T592A and T592E mutations into the endogenous *SAMHD1* locus of BlaER1 cells.

**Figure 3:**
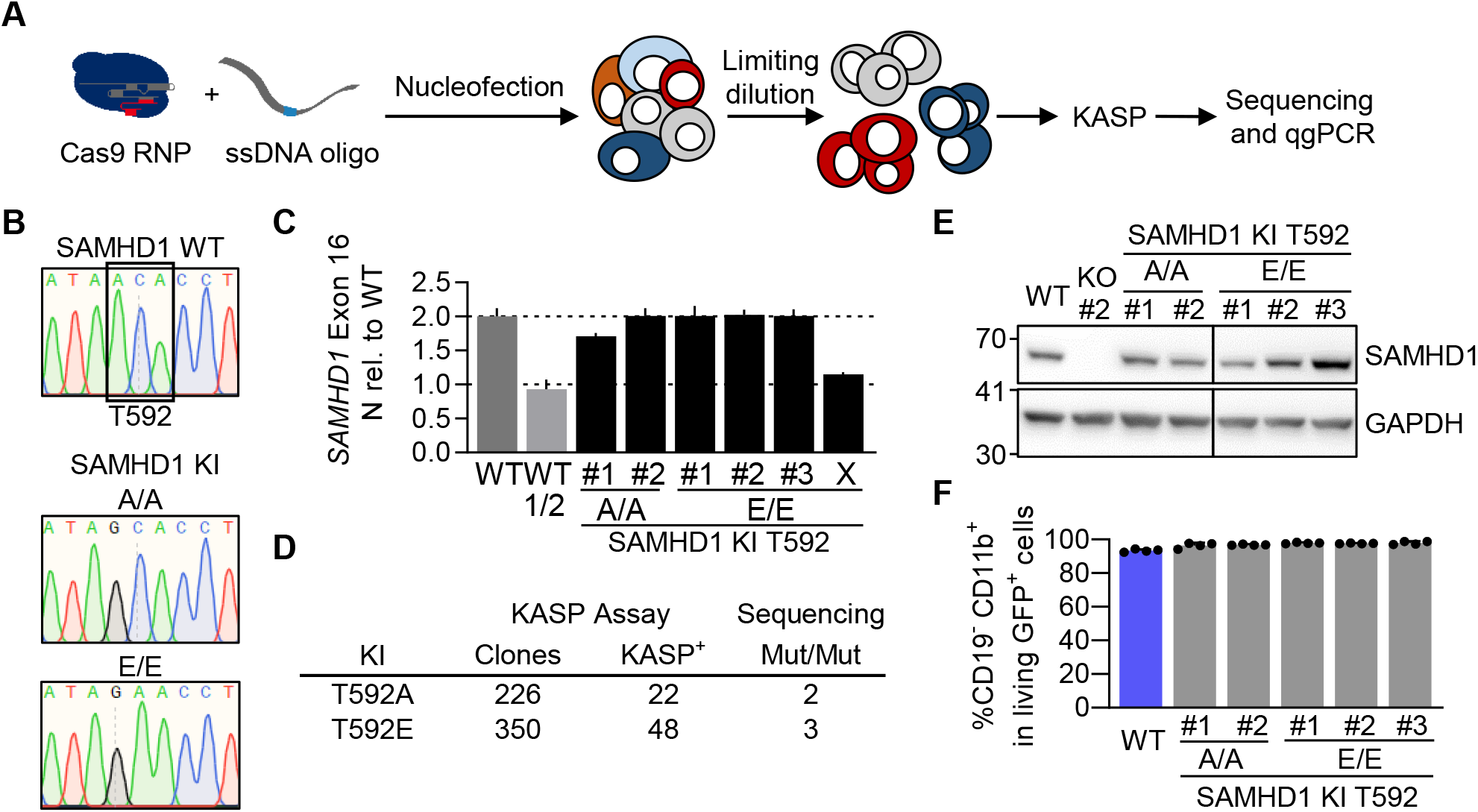
A pipeline to generate mutants of SAMHD1 by CRISPR/Cas9 mediated knock-in. **(A)** Schematic representation of CRISPR/Cas9 meditated knock-in (KI) to generate mutants of SAMHD1 in BlaER1 cells. Cas9 ribonucleoprotein (RNP) together with ssDNA correction template was introduced into BlaER1 cells via nucleofection. Single cell clones generated by limiting dilution were screened using KASP assay and validated by Sanger sequencing and quantitative genomic PCR (qgPCR) (30). **(B)** Representative sections of Sanger sequencing traces obtained from genomic *SAMHD1* exon 16, highlighting successful mono- or bi-allelic single base exchange at the base triplet corresponding to amino acid position T592 in BlaER1 SAMHD1 KI T592A and T592E mutant single cell clones. No further mismatches were detected up- or downstream of shown section in the amplified region. Two independent sequencing runs were performed. Homozygous T592E mutants were additionally confirmed by allele specific sequencing after TA-cloning. **(C)** Quantitative genomic PCR for *SAMHD1* exon 16 against reference gene *TERT* was performed and 2^−Δct^ value obtained from SAMHD1 KI clones normalized to WT in order to obtain the allele number. As a control half of the WT (WT 1/2) DNA was inoculated and Δct of *SAMHD1* was calculated against ct of *TERT*, which was obtained in the WT with normal DNA amount. Error bars indicate standard deviation of technical triplicates in a representative experiment (n = 2). **(D)** Number of single cell clones obtained from CRISPR/Cas9 RNP and ssDNA correction oligo treated BlaER1 cells and number of clones scoring positive in KASP assay, as well as homo- (Mut/Mut) mutants identified by Sanger sequencing and confirmed by qgPCR are shown. **(E)** Transdifferentiated SAMHD1 KI BlaER1 cells were analyzed by immunoblot for SAMHD1 expression and compared to WT cells. GAPDH was used as a loading control (representative for n = 3). **(F)** Percentage of CD19^−^ CD11b^+^ cells in living GFP^+^ transdifferentiated WT and SAMHD1 KI cells were quantified by flow cytometry. Error bars indicated standard deviation of biological replicates (n = 4).

### Homozygous SAMHD1 T592E mutation increases HIV-1 infection in transdifferentiated BlaER1 cells

We infected several clones of transdifferentiated homozygous SAMHD1 phosphoablative T592A and phosphomimetic T592E KI BlaER1 cell mutants with HIV-1-mCherry reporter virus and measured the fold change of %mCherry^+^ cells in CD19^+^ living GFP^+^ cells relative to infection in WT cells. In all three clones, homozygous SAMHD1 T592E mutation significantly increased HIV-1-mCherry infection up to 31-fold (T592E/T592E Clone #1 *p = 0.0017*, #2 and #3 *p < 0.0001*, n = 3, One-way ANOWA) (Fig. 4A and 4B). In contrast, SAMHD1 T592A KI mutants completely retained their restrictive potential in transdifferentiated BlaER1 cell clones and behaved similar to WT BlaER1 cells upon challenge with HIV-1-mCherry (Fig. 4A and 4B). Using CRISPR/Cas9 KI, we were able to validate the loss of HIV-1 restriction in SAMHD1 phosphomimetic T592E mutants in macrophage-like cells. In this model, mutants of SAMHD1 are analyzed in the native genomic context and show physiological expression levels, confirming the role of T592 phosphorylation in the regulation of the anti-viral activity of SAMHD1.

**Figure 4:**
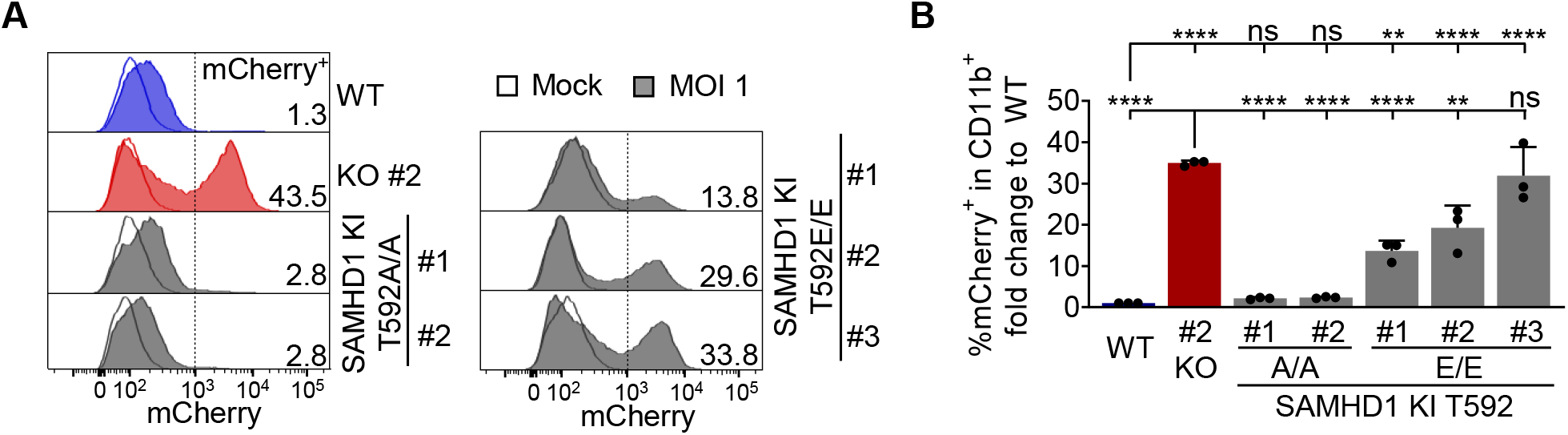
Homo- and heterozygous SAMHD1 T592E, but not T592A mutation leads to loss of HIV-1 restriction in transdifferentiated BLaER1 cells. **(A, B)** Transdifferentiated homozygous SAMHD1 T592E and T592A BlaER1 KI clones were infected with VSV-G pseudotyped HIV-1 single cycle mCherry reporter virus pNL4.3 IRES mcherry E^−^ R^+^ at MOI 1. Percentage of mCherry^+^ cells was quantified by flow cytometry in living GFP^+^ CD11b^+^ BlaER1 cells at 24 hpi. **(A)** Representative histograms are shown for mock and HIV-1 mCherry reporter virus infected cells. Percentage of mCherry^+^ cells in living GFP^+^ CD11b^+^ BlaER1 cells is indicated (n = 3). **(B)** To calculate fold change, percentage of mCherry^+^ cells in infected SAMHD1 KI clones was normalized to WT. Bar graphs indicate mean of experiments, dots individual biological replicates. Error bars correspond to standard deviation (n = 3, One-way ANOWA, ** p < 0.01; **** p < 0.0001; ns, not significant).

### SAMHD1 T592E or T592A knock-in does not affect dNTP levels in transdifferentiated BlaER1 cells

Previous reports on the effect of SAMHD1 T592 phosphorylation on SAMHD1 dNTPase activity were in-conclusive (1). In order to correlate HIV-1 restrictive potential in transdifferentiated BlaER1 cells with cellular dNTP pool size and thus SAMHD1 dNTPase activity, we measured intracellular dNTP levels by primer extension assay. Transdifferentiated WT BlaER1 cells contained low amounts of dATP (846 ± 63 fmol/10^6^ cells, n = 5), dCTP (788 ± 117 fmol/10^6^ cells, n = 5), dGTP (724 ± 94 fmol/10^6^ cells, n = 5) and dTTP (933 ± 342 fmol/10^6^ cells, n = 5). Depletion of the minor fraction of CD19^+^ cells after transdifferentiation further reduced the levels of dATP (578 fmol/10^6^ cells), dCTP (661 fmol/10^6^ cells), dGTP (295 fmol/10^6^ cells) and dTTP (448 fmol/10^6^ cells). Since activity of HIV-1 RT is likely to be dependent on cellular dNTP concentrations rather than total dNTP pools, we determined cellular dNTP concentrations as a function of transdifferentiated BlaER1 cell volumes (569 ± 138 μm^3^, n_cells_ = 15). We found transdifferentiated WT BlaER1 cells to harbor dNTP concentrations (Tab. 1), similar or lower to those found in resting T cells (31). Depletion of incompletely transdifferentiated (CD19+) cells from bulk preparations of transdifferentiated BlaER1 cells further reduced dNTP concentrations (Tab. 1). As expected, SAMHD1 KO led to a significant increase in cellular dATP (2.3-fold, *p < 0.0001*, One-way ANOWA), dGTP (3.2-fold, *p < 0.0001*, One-way ANOWA) and dTTP (2.2-fold, *p < 0.0001*, One-way ANOWA) levels in transdifferentiated BlaER1 cells, as compared to WT cells (Fig. 5A). In contrast, neither homozygous SAMHD1 T592E, nor T592A mutations led to an increase of cellular dNTP levels (Fig. 5A). Since SAMHD1 KO only slightly affected cellular dCTP levels, dNTP composition in transdifferentiated BlaER1 SAMHD1 KO cells was altered. In contrast, neither SAMHD1 T592E nor SAMHD1 T592A KI mutants showed consistent differences in cellular dNTP composition (Fig. 5B). In summary, dNTP measurements in transdifferentiated BlaER1 cells, harboring homozygous phosphomimetic T592E or phosphoablative T592A mutations in the endogenous *SAMHD1* locus indicate that phosphorylation at SAMHD1 residue T592 has no impact on cellular dNTP pools and is therefore unlikely to regulate SAMHD1 dNTPase activity in cells.

**Figure 5:**
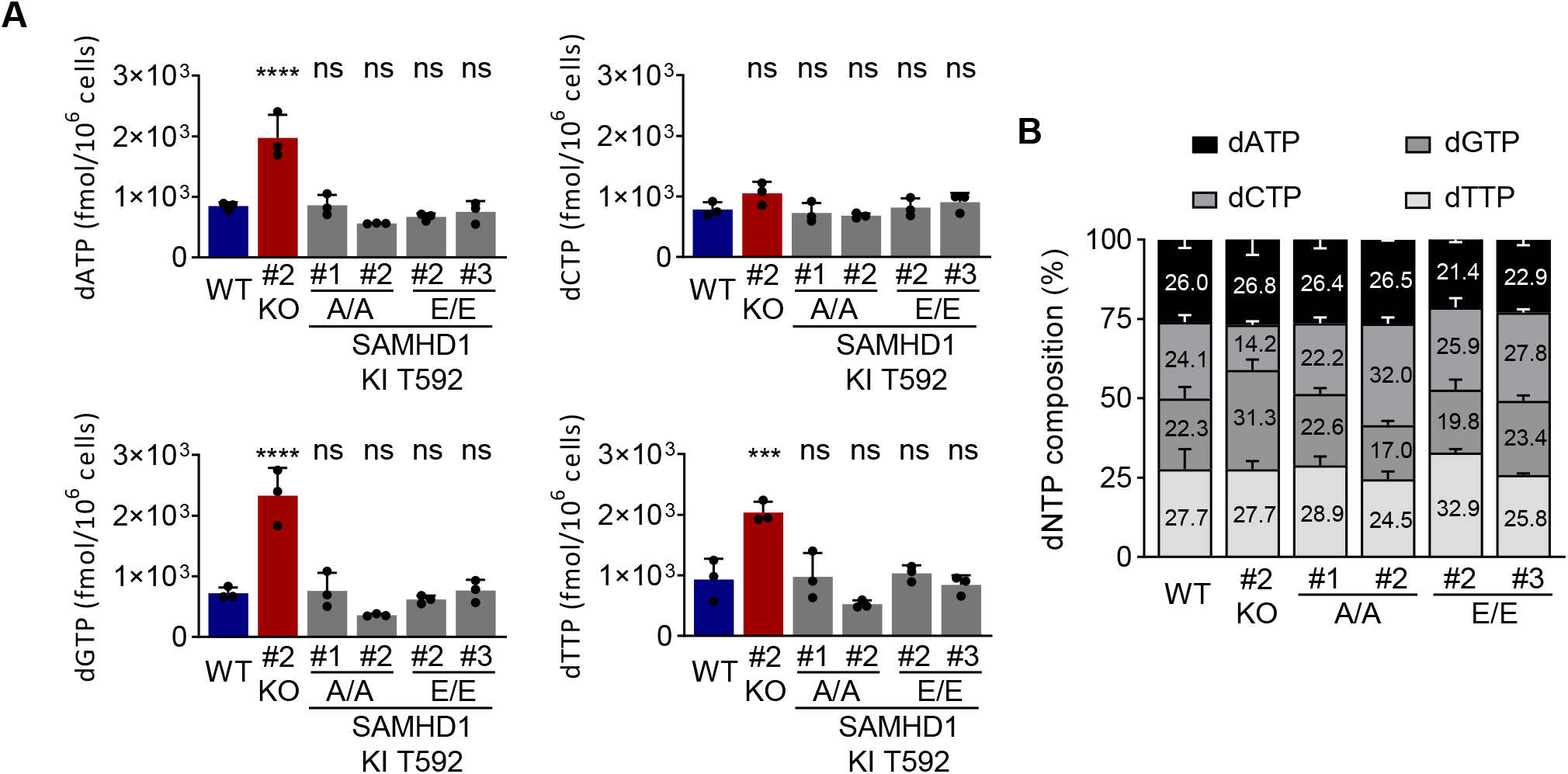
SAMHD1 T592E or T592A knock-in does not affect dNTP levels in transdifferentiated BlaER1 cells. **(A, B)** Cellular dNTP levels were measured in transdifferentiated homozygous SAMHD1 T592E and T592A BlaER1 KI mutants. dNTP amounts were compared to transdifferentiated WT BlaER1 cells. **(A)** Amount of indicated dNTP is depicted per 1 × 10^6^ cells. Bar graphs indicate mean of experiments, dots individual biological replicates. Error bars correspond to standard deviation (n = 3, One-way ANOWA, *** p < 0.01; **** p < 0.0001; ns, not significant). **(B)** dNTP composition in individual BlaER1 SAMHD1 KI clones is shown, with total dNTP content set as 100%. Error bars indicate standard deviation (n = 3).

**Table 1:** dNTP concentrations in transdifferentiated and CD19 depleted BlaER1 cells.

| Cellular concentration (μM) | dATP | dCTP | dGTP | dTTP | n |
| --- | --- | --- | --- | --- | --- |
| WT BLAER1 cells (td) | 1.51 | 1.22 | 1.40 | 1.83 | 5 |
| SAMHD1 KO BLAER1 cells (td) | 4.39 | 1.89 | 5.92 | 5.88 | 5 |
| WT CD19<sup>-</sup> BLAER1 cells (td) | 1.02 | 1.16 | 0.52 | 0.79 | 1 |
| SAMHD1 KO CD19<sup>-</sup> BLAER1 cells (td) | 2.21 | 1.54 | 2.31 | 2.03 | 1 |
| Resting T cells (31) | 1.72 | 1.88 | 1.51 | 1.67 | 3 |
| Activated T cells (31) | 5.09 | 5.91 | 4.53 | 7.91 | 3 |

## Discussion

SAMHD1 is a major cellular dNTPase and a potent HIV-1 restriction factor (2, 4, 3, 5, 6, 10, 32). However, whether SAMHD1 dNTPase activity is meditating its anti-viral activity and how this is regulated by T592 phosphorylation is still a matter of debate (1, 14).

To tackle this question with a novel toolkit, we used transdifferentiated BlaER1 cells as an alternative and versatile myeloid model to study HIV-1 infection in macrophage-like cells. Transdifferentiated BlaER1 cells closely resemble human macrophages and have successfully been used to study innate immune signaling (28, 29). In contrast to PMA activated THP-1 or U937 cells, BlaER1 cell transdifferentiation relies on the activation of the fusion protein of the macrophage transcription factor C/EBPα and the estradiol receptor leading to a switch of a myeloid cell program (29, 33). This cell system is an interesting physiological model for HIV-1 infection in macrophage-like cells, as we were able to show that SAMHD1 is completely dephosphorylated at residue T592 in transdifferentiated BlaER1 cells and serves as a major restriction factor for HIV-1.

To test mutants of SAMHD1 for anti-viral restriction activity in a physiological genetic and relevant cellular context, we combined transdifferentiated BlaER1 cells, as a novel myeloid model, with CRISPR/Cas9 mediated knock-in. We developed a gene editing strategy using CRISPR/Cas9 RNP and ssDNA oligo introduction by nucleofection, KASP screening of single cell clones and rigorous validation by sequencing and qgPCR (Fig. 3A). By introducing point mutations that correspond to phosphoablative T592A and phosphomimetic T592E mutations into the genomic *SAMHD1* locus, we were able to genetically uncouple SAMHD1 mediated anti-viral restriction and cellular dNTPase activity. For the first time, we were able to validate the phenotypic consequence of SAMHD1 T592E phosphomimetic mutation, and hence the effect of T592 phosphorylation, for its anti-viral restriction activity in a myeloid model that is not based on overexpression.

Overexpression of SAMHD1 mutants in U937 background using retroviral transduction has several technical limitations. Even though PMA-activated U937 cells are often considered not to express SAMHD1, they actually can express small amounts of endogenous SAMHD1, which can be further enhanced upon interferon treatment (34). Presence of endogenous WT SAMHD1 might affect the function of overexpressed mutant SAMHD1, especially if heterotetramers are formed. In addition, strong exogenous, often viral, promotors drive SAMHD1 expression here, leading to high expression levels and to non-physiological phosphorylation ratios, *i.e.* hyperphosphorylation (data not shown). In mutants generated by CRISPR/Cas9 KI, modified SAMHD1 is expressed in the physiological genomic context from the endogenous promotor and thus under normal transcriptional regulation. In BlaER1 cells, SAMHD1 expression is very low in native cells, but strongly induced upon transdifferentiation (Fig. 1C). This is also the case for SAMHD1 mutants. The use of CRISPR/Cas9 KIs avoids potential effects of constitutively expressed SAMHD1 mutants on cycling BlaER1 cells.

In transdifferentiated WT BlaER1 cells, we measured dNTP levels and concentrations which were similar or slightly lower than those found in resting T cells (Tab. 1) (31). After depletion of CD19^+^ incompletely transdifferentiated cells from bulk preparations of transdifferentiated BlaER1 cells, we were able to further reduce the levels of all dNTPs (Tab. 1) (31). Considering HIV-1 reverse transcriptase Km and Kd values measured *in vitro*, this indicates that the dNTP concentrations found in transdifferentiated BlaER1 cells could in principle be low enough to restrict or delay HIV-1 RT (35–38).

Concomitantly, SAMHD1 KO increased cellular dNTP concentrations in transdifferentiated BlaER1 cells up to 4-fold (Fig. 5A), which is reminiscent of the 5- to 8-fold increase upon T cell activation (31). In stark contrast however, neither endogenous SAMHD1 T592E nor T592A mutation increased cellular dNTP concentrations in transdifferentiated BlaER1 cells (Fig. 5A). This indicates that the loss of restriction observed in endogenous T592E mutants is probably not caused by increased dNTP levels or reduced SAMHD1 dNTPase activity in transdifferentiated BlaER1 cells. In addition, SAMHD1 T592E and T592A mutations had no consistent effect on dNTP pool composition in transdifferentiated BlaER1 cells (Fig. 5B), ruling out an effect of the phosphomimetic mutation on SAMHD1 dNTPase substrate preferences and thus dNTP ratios, as proposed earlier (39). More specifically, endogenous SAMHD1 T592E mutations did not increase cellular dCTP concentration (Fig. 5A). Taken together mutagenic analysis of SAMHD1 residue T592 indicates that SAMHD1 dNTPase activity or substrate preference in transdifferentiated BlaER1 cells is not regulated by phosphorylation at this specific residue. Consequently, loss of HIV-1 restriction in SAMHD1 T592E mutants cannot be attributed to changes in SAMHD1 dNTPase activity. Thus, in line with previous reports, our data confirm that SAMHD1 anti-viral activity is not, or at least not exclusively, mediated by SAMHD1 dNTPase activity (14). SAMHD1 T592 phosphorylation might regulate additional functional entities of SAMHD1, which could be necessary for full anti-viral restriction capacity. A deeper investigation is needed to understand the functional cause of restriction in relevant HIV-1 target cells.

SAMHD1 regulation certainly is certainly more complex than commonly assumed. In addition to multiple potential phosphorylation sites, SAMHD1 is modified by acetylation, SUMOylation, ubiquitination and O-GlcNAcylation (20, 21, 40–47). Also, SAMHD1 was demonstrated to harbor redox sensitive cysteine residues, which affect both dNTPase and anti-viral activity (48, 49). SAMHD1 acetylation at residue K405 was shown to affect SAMHD1 dNTPase activity *in vitro.* However, the effect on its anti-viral restriction activity is unclear (42). O-GlcNAcylation at S93 was proposed to increase SAMHD1 stability and thus hepatitis B virus restriction (47). Recently, SAMHD1 SUMOylation at residue K595 was shown to be required for HIV-1 restriction in PMA differentiated U937 cells. PMA differentiated U937 cells in which SAMHD1 mutants that abrogate restriction and SUMOylation at residue K595 were overexpressed did not show increased dATP levels, phenocopying phosphomimetic T592 mutants of SAMHD1 (50). It will be interesting to investigate in more detail how T592 phosphorylation and K595 SUMOylation are integrated and it will be crucial to validate SAMHD1 (co-) regulation via divers proposed post-translational modifications in physiological settings. Post-translational regulation of SAMHD1 might not only be achieved by the direct modification of single residues, but also by interaction partners, that could modulate or mediate SAMHD1 anti-viral activity.

A better understanding SAMHD1 regulation in relevant HIV-1 target cells, will also improve our understanding of how SAMHD1 inhibits HIV-1 replication and which conditions license SAMHD1 anti-viral capacity.

## Material and Methods

### Cell lines

Human 293T/17 (ATCC No.: CRL-11268) cells were cultured in DMEM (Sigma-Aldrich) supplemented with 10% fetal calf serum (FCS; Sigma-Aldrich) and 2 mM L-glutamine (Sigma-Aldrich) at 37°C and 5% CO_2_. Human BlaER1 cells (a kind gift of Thomas Graf) (29) cells were grown in RPMI (Sigma-Aldrich) supplemented with 10% FCS and 2 mM L-glutamine at 37°C and 5% CO_2_. For transdifferentiation, 1 × 10^6^ BlaER1 cells per well of a 6-well tissue culture plate were treated with 10 ng/ml human recombinant M-CSF and IL-3 (PeproTech) and 100 nM β-estradiol (Sigma-Aldrich) for 7 days. Half of the cell culture supernatant was replaced with medium containing cytokines and β-estradiol at days 2 and 6. All cell lines were free of mycoplasma contamination, as tested by PCR Mycoplasma Test Kit II (PanReac AppliChem).

### CRISPR/Cas9 knock-out and knock-in

For CRISPR/Cas9 mediated SAMHD1 knock-out (KO), 200 pmol Edit-R Modified Synthetic crRNA targeting *SAMHD1* exon 1 (crSAMHD1_ex1, target sequence: 5’-ATC GCA ACG GGG ACG CTT GG, Dharmacon), 200 pmol Edit-R CRISPR-Cas9 Synthetic tracrRNA (Dharmacon) and 40 pmol Cas9-NLS (QB3 Macrolab) were assembled *in-vitro*, as previously described (51). Ribonucleoproteins were introduced into 1 × 10^6^ sub-confluent BlaER1 cells using 4D-Nucleofector X Unit and SF Cell line Kit (Lonza), applying program DN-100. Single cell clones were generated using limited dilution one day after nucleofection. To confirm bi-allelic SAMHD1 KO, the modified region was amplified using primer SAM_Seq_Gen-23_FW (5’-GAT TTG AGG ACG ACT GGA CTG C) and SAM_Seq_Gen1116_RV (5’-GTC AAC TGA ACA ACC CCA AGG T) together with GoTaq polymerase (Promega), followed by cloning into pGEM T-easy vector system (Promega) and Sanger sequencing. For knock-in (KI), 100 pmol of the respective ssDNA homologous recombination template with 30 bp homology arms to introduce T592A (5’-TAG GAT GGC GAT GTT ATA GCC CCA CTC ATA GCA CCT CAA AAA AAG GAA TGG AAC GAC AGT A, Dharmacon) or T592E (5’-TAG GAT GGC GAT GTT ATA GCC CCA CTC ATA GAA CCT CAA AAA AAG GAA TGG AAC GAC AGT AC) was nucleofected together with ribonucleoprotein complex containing crSAMHD1_ex16 (target sequence: 5’-TTT TTT TGA GGT GTT ATG AG, Dharmacon). When single cell clones reached confluency, duplicates were generated. One half was lysed (10 min, 65°C; 15 min, 95°C) in lysis buffer (0.2 mg/ml Proteinase K, 1 mM CaCl_2_, 3 mM MgCl_2_, 1 mM EDTA, 1% Triton X-100, 10 mM Tris (pH 7.5)) (52) and screened for successful KI using mutation specific custom designed KASP genotyping assays (LGC) and KASP V4.0 2x Master mix (LGC) on a CFX384 Touch Real-Time PCR Detection System (BioRad). Homozygous KI was confirmed by Sanger sequencing after amplification using primer SAM_Seq_Gen58570_FW (5’-CAT GAA GGC TCT TCC TGC GTA A) and SAM_Seq_Gen59708_RV (5’-ACA AGA GGC GGC TTT ATG TTC C) together with KOD Hot Start DNA Polymerase (Merck). Additionally, allele specific sequencing as described for SAMHD1 KO was performed, if required. Presence of large deletions in the region between amplification primers was excluded by PCR and analytic gel electrophoresis. Presence of both alleles was confirmed by quantitative genomic PCR (30), performed using *SAMHD1* exon 16 specific PrimeTime qPCR Assay (FW: 5’-CTG GAT TGA GGA CAG CTA GAA G, RV: 5’-CAG CAT GCG TGT ACA TTC AAA, Probe: /56-FAM/ AAA TCC AAC /Zen/ TCG CCT CCG AGA AGC /3IABkFQ/, IDT), human TERT TaqMan Copy Number Reference (Thermo Fischer) and PrimeTime Gene Expression Master Mix (IDT) on a CFX384 machine.

### HIV-1 reporter virus infection

VSV-G pseudotyped HIV-1 reporter viruses pNL4.3 E^−^ R^−^ luc (53) (HIV-1-luc) and pNL4.3 IRES mCherry E^−^ R^+^ (HIV-1-mCherry) were produced, as detailed previously (22). Briefly, pNL4.3 E^−^ R^−^ luc (a kind gift of Nathaniel Landau) or pNL4.3 IRES mCherry E^−^ R^+^ (a kind gift of Frank Kirchhoff) were co-transfected together with pCMV-VSV-G into 293T/17 cells using 18 mM polyethylenimine (Sigma-Aldrich). Filtered (0.45 μm) supernatants were treated with 1 U/ml DNAse I (NEB; 1 h, RT) and purified through a 20% sucrose cushion (2 h, 106750g, 4°C). Viral stocks were titrated for β-galactosidase activity on TZM-bl cells. Virus-like particles containing Vpx (VLP-Vpx) were produced in an analogue manner using pSIV3+ (54) derived from SIVmac251 (a kind gift of Nicolas Manel) and pCMV-VSV-G. The amount of VLP-Vpx used in all experiments was optimized for complete SAMHD1 degradation. For infection 3 × 10^4^ cells were seeded per well of a 96-well tissue culture plate. Transdifferentiated BlaER1 cells were allowed to settle for 2 h in medium without cytokines and β-estradiol. VSV-G pseudotyped HIV-1 reporter virus at indicated MOI, as well as VLP-Vpx, were added, followed by spin occulation (1.5 h, 200g, 32°C). Infection was quantified after 24 h by FACS (for HIV-1-mCherry) or by adding 50 μl/well britelite plus reagent (PerkinElmer) and measurement on a Pherastar FS (BMG) (for HIV-1-luc). To show VLP-Vpx mediated SAMHD1 degradation, 4.4 × 10^5^ transdifferentiated BlaER1 cells were treated in a 12-well tissue culture plate in the same conditions and concentrations as stated above.

### Flow Cytometry

For FACS analysis of BlaER1 transdifferentiation, 1 × 10^6^ native or transdifferentiated BlaER1 cells were collected, washed once in FACS buffer (10% FCS, 0.1% Sodium acetate in PBS; 10 min, 300g, 4°C) and stained with CD11b-APC (M1/70, Biolegend), CD19-PE (HIB19, Biolegend) or respective isotype controls (Biolegend) and Fixable Viability Dye eFluor 780 (Thermo Fischer) in presence of FC Block (BD, 20 min, 4°C). Stained cells were washed in FACS buffer twice and fixed in 2% paraformaldehyde (30 min, RT), before analyzing on a LSR II instrument (BD). For readout of HIV-1-mCherry infection, six wells of a 96-well plate were pooled and stained with CD11b-APC and Fixable Viability Dye eFluor 780 as detailed above. Infected cells were analyzed on a BD LSRFortessa.

### Immunoblot

For immunoblot, cells were washed in PBS, lysed in radioimmunoprecipitation buffer (RIPA; 2 mM EDTA, 1% glycerol, 137 mM NaCl, 1% NP40, 0.1% SDS, 0.5% sodium deoxycholate, 25 mM Tris (pH 8.0)) supplemented with proteinase and phosphatase inhibitor (Roche) for 30 min on ice. Lysate was cleared (30 min, 15000g, 4°C) and protein content was measured by Bradford assay using Protein Assay Dye Reagent Concentrate (BioRad). 20 μg total protein were denatured (10 min, 70°C) in NuPAGE LDS Sample Buffer and Reducing Reagent (Thermo Fischer) and separated on a NuPAGE 4-12% Bis-Tris gradient gel (Thermo Fischer) in MOPS running buffer (1 M MOPS, 1 M Tris, 69.3 mM SDS, 20.5 mM EDTA Titriplex II). Transfer was performed in an XCell II Blot Module in NuPAGE Transfer Buffer (Thermo Fischer) onto a Hybond P 0.45 PVDF membrane (GE Healthcare). After blocking in 5% BSA or milk powder (Carl Roth) in TBST (Tris-buffered saline, 0.1% Tween; 2 h, 4°C), primary antibodies anti-GAPDH (14C10, CST), anti-Cyclin B1 (4138, CST), anti-Cyclin A2 (4656, CST), anti-SAMHD1 (12586-1-AP, Proteintech), anti-SAMHD1 (A303-691A, Bethyl) and anti-SAMHD1-pT592 (D702M, CST) diluted in 5% BSA or milk powder in TBST were applied overnight at 4°C. Subsequent to washing in TBST, anti-rabbit IgG, horseradish peroxidase (HRP)-linked antibody (CST) was applied (2 h, 4°C) and the membrane was washed again before detection on a FUSION FX7 (Vilber Lourmat) using ECL Prime reagent (GE). If required, membranes were stripped of bound antibody in stripping buffer (2% SDS, 62.5 mM Tris-HCl (pH 6.8), 100 mM β-mercaptoethanol; 1 h, 65°C). Band densities were determined with FUSION software (Vilber Lourmat).

### Cellular dNTP levels and concentrations

For measurement of cellular dNTP levels, 2 × 10^6^ transdifferentiated BlaER1 cells were washed in PBS and subjected to methanol extraction of dNTPs, followed by quantification of all four dNTPs by single nucleotide incorporation assay, as described previously (31). CD19 depletion was performed using CD19 microbeads and MS columns (Miltenyi). Cell volumes were determined by seeding respective cell types on a Poly-D-Lysine (Sigma) coated (10%, 1.5h, RT) Cell Carrier-96 well plate (Perkin Elmer). After centrifugation (5 min, 300g), cells were fixed (4% PFA, 15 min, 37°C), permeabilized (0.1% Triton X-100, 5 min, 37°C) and stained using HCS CellMask Deep Red Stain (Thermo Fischer, 30 min, RT). Z-Stack of stained cells was acquired using confocal imaging platform Operetta (Perkin Elmer) and volume was calculated as a sum of cell areas in all relevant Z-stacks using Harmony software (Perkin Elmer).

### Statistical analysis

Statistical analysis was performed using GraphPad Prism (V8). Mean and standard deviations are shown. Statistical significance was assessed using unpaired two-tailed t-test, as well as non-parametric Kruskal-Wallis test or parametric One-Way ANOWA, corrected against multiple testing using Dunn’s or Dunnet correction, respectively.

## Acknowledgements

The authors thank Michaela Neuenkirch and Saskia Mönch for technical assistance, as well as Thomas Graf (Centre for Genomic Regulation Barcelona, Spain) for the BlaER1 cell line and Stefan Bauer (University of Marburg, Germany) for providing BlaER1 cell transdifferentiation protocols. In addition, the authors thank Nathaniel R. Landau (NYU School of Medicine, USA) for pNL4.3 E^−^ R^−^ luc, Frank Kirchhoff (University of Ulm, Germany) for pNL4.3 IRES mCherry E^−^ R^+^ and Nicolas Manel (Institut Curie, Paris) for pSIV3^+^ plasmid.

## Author Contributions

Conceptualization, M.S., R.K. methodology and formal analysis, M.S., R.K., A.O., N.H.F.; investigation, M.S., P.R., K.S., A.O., N.H.F.; resources, B.K., R.K.; writing—original draft preparation, M.S., R.K.; writing—review and editing, R.K., M.S.; visualization, M.S.; supervision, R.K.; funding acquisition, R.K., B.K.; All authors have read and agreed to the published version of the manuscript.

